# *In planta genome* editing with CRISPR/Cas9 ribonucleoproteins

**DOI:** 10.1101/2021.06.24.449705

**Authors:** Yuya Kumagai, Yuelin Liu, Haruyasu Hamada, Weifeng Luo, Jianghui Zhu, Misa Kuroki, Yozo Nagira, Naoaki Taoka, Etsuko Kato, Ryozo Imai

**Affiliations:** Genome Edited Crop Development Group, Division of Crop Genome Editing, Institute of Agrobiological Sciences, National Agriculture and Food Research Organization (NARO), Kannondai 3-1-3, Tsukuba, Ibaraki, 3058602, Japan; Agri-Bio Research Center, Biotechnology Research Laboratories, KANEKA CORPORATION, Iwata, Shizuoka, 438-0802, Japan; Structural Biology Team, Advanced Analysis Center, National Agriculture and Food Research Organization, Tsukuba, Ibaraki 305-8602, Japan

**Author notes:** Corresponding author: Ryozo Imai. Institute of Plant and Food Science, Department of Biology, Southern University of Science and Technology, Shenzhen, Guangdong 518055, China (Yuelin Liu).

## Abstract

*In planta* genome editing represents an attractive approach to engineering crops/varieties that are recalcitrant to culture-based transformation methods. Here, we report the direct delivery of CRISPR/Cas9 ribonucleoproteins into the shoot apical meristem using *in planta* particle bombardment and introduction of a *semidwarf1 (sd1)*-orthologous mutation into wheat. The triple knockout *tasd1* mutant of an elite wheat variety reduced culm length by 10% without a reduction in yield.

## Main

Shoot apical meristems (SAMs) maintain the potential to develop into floral organs. SAMs are generally composed of three independently dividing cell layers (L1 to L3), among which the cells of the sub-epidermal layer (L2) are destined to develop into germ cells, such as pollen grains and embryo sacs ^1,2^. We developed a new transformation method, utilizing *in planta* particle bombardment (iPB), in which SAM tissue of wheat (*Triticum aestivum*) is subjected to biolistic transformation^3^. As this method does not require pre- or post-transformation tissue culture methodologies, it can be used to transform recalcitrant cultivars. With this method, transiently-expressed, clustered regularly interspaced short palindromic repeat (CRISPR)/Cas9 has been used to create genome edited plants^4^. Efficient DNA-free genome-editing systems have been recently developed using CRISPR/Cas9 ribonucleoproteins (RNPs) in plants. Direct delivery of CRISPR/Cas9 RNP into protoplasts^5^, fertilized eggs^6^, or immature embryos^7,8^ has been successfully used to create genome-edited plants. These methods, however, require callus tissue culture and regeneration steps which may limit the application of this approach strictly to varieties that are amenable to cell/tissue culture. Here, we developed a direct delivery system of Cas9/gRNA RNP into SAMs and established a non-culture method to solve this limitation.

As shown in Fig.1a, we delivered gold particles coated with CRISPR/Cas9 RNPs to wheat SAMs (iPB-RNP method) as previously described^3,4^ and screened E_0_ genome-edited mutants by a cleaved amplified polymorphic sequences (CAPS) assay with 5^th^ leaves. We first used *TaQsd1* as a target site and identified five E_0_ positive mutants after two rounds of screening with a CAPS assay (Fig.1b) which were subsequently validated by Sanger sequencing (Fig.1c). A particular plant (Q2) was identified to contain mutations in all three homoeologous genes (Fig.1c). In addition to screening the *TaQsd1* locus, we also deployed this strategy with additional target sites (TaOr_t0, TaOr_t1, TaHRGP-like1_t2, Supplementary Table 3) and obtained promising editing efficiency (from 1% to 8.3%) in E_0_ plants (Fig. 1d). Collectively, these results demonstrated that the iPB-RNP method is capable of being deployed for *in planta* genome editing with comparable efficiency to the iPB-DNA method^9^.

**Figure 1.**
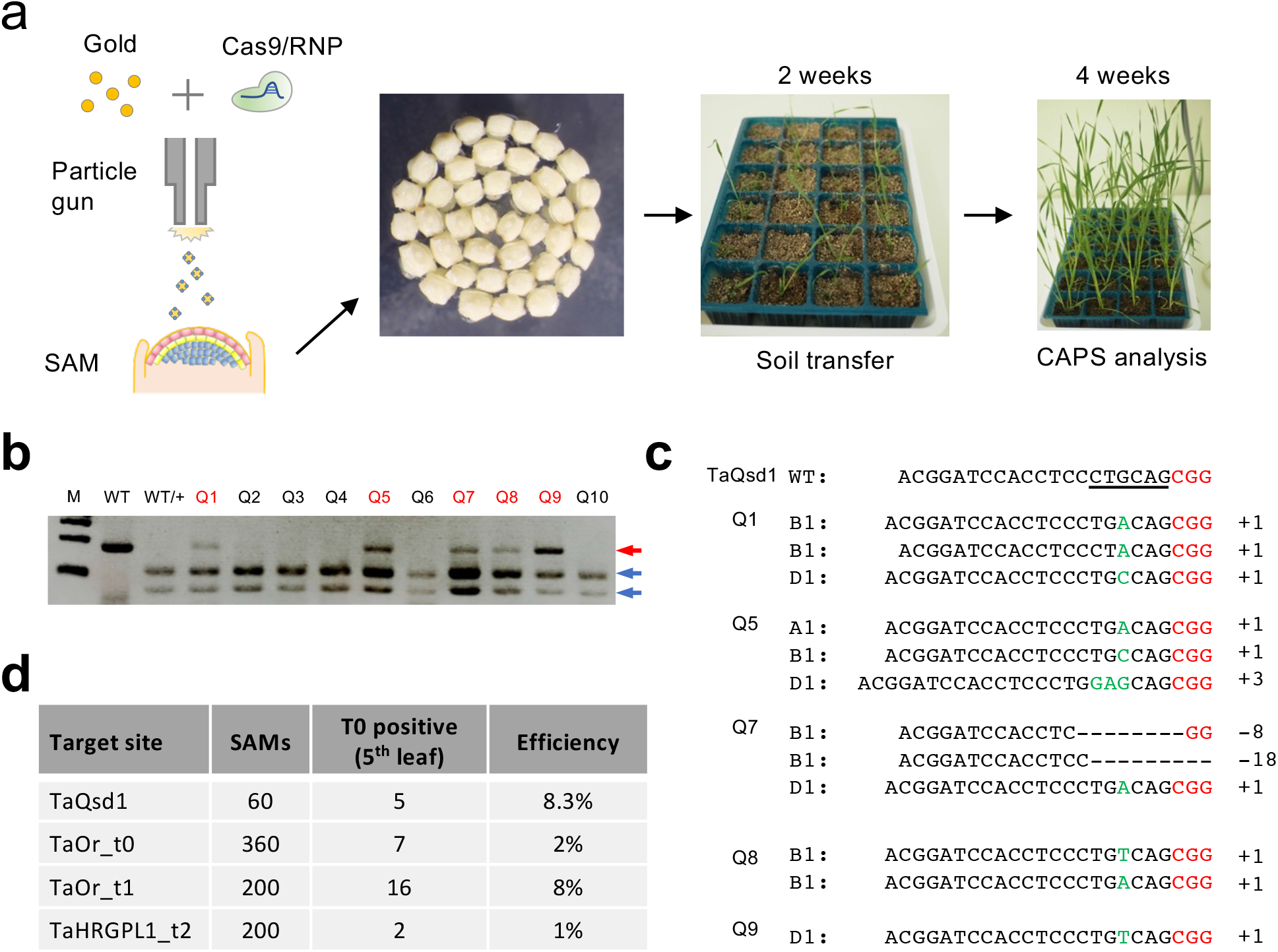
*In planta* RNP-mediated genome editing in wheat. **a**, The workflow of iPB-RNP method utilizing wheat SAMs. **b**, CAPS analysis of E_0_ plants at the *TaQsd1* locus. The PCR products were amplified by an A, B, and D genome common primer set. WT, undigested PCR products; WT/+, *Pst* I digested PCR products. Red and blue arrows indicate undigested and digested bands after *Pst* I treatment, respectively. A 100 bp ladder was used as a size marker. **c**, The genotypes of Q1, Q5, Q7, Q8 and Q9 as identified by sequencing. The black and red characters indicate the gRNA and PAM sequences, respectively. The *Pst* I restriction site is underlined. Nucleotides inserted are shown in green characters. **d**, Summary of genome editing experiment on locus sites of TaQsd1, TaOr_t0, TaOr_t1 and TaHRGP-like_t2 using the iPB-RNP method.

Currently, most wheat commercial cultivars carry a dominant allele of *REDUCED HEIGHT 1* (*Rht1*), a “Green Revolution” gene, encoding a GAI/DELLA protein^10,11^, and express a semidwarf phenotype due to partial gibberellic acid (GA) insensitivity. In contrast, the rice (*Oryza sativa*) semidwarf gene (*sd1, semidwarf 1*) encodes a GA20 oxidase, which is involved in GA biosynthesis^12,13,14^. The impact of the dominant/GA-insensitive and recessive/GA-deficient alleles in wheat and rice, respectively, are affected by their ploidy level. Using genome editing strategies, it is plausible to introduce recessive *sd1* mutation in *Rht1* wheat and evaluate the effect of the double mutation. With a BLAST search of the Gramene database (http://www.gramene.org), we identified three homoeologous genes, TraesCS3A02G406200, TraesCS3B02G439900, and TraesCS3D02G401400, which encode proteins with 77-78% identity to the rice *sd1* gene (*OsGA20ox2*). A phylogenetic tree of rice and wheat GA20 oxidases identified four clades, each of which contains one rice and three or four wheat homoeologous genes (Supplementary Fig. 1). These results suggest that GA20 oxidases within a clade have an evolutionary conserved function. Thus, we concluded that *TaSD-A1, TaSD-B1* and *TaSD-D1* were the three wheat orthologs representing homoeologous genes to rice *sd1*.

To create a *tasd1* triple knockout mutant using CRISPR/Cas9 RNP, three single-guide RNA (sgRNA) target sequences (target_1, _2, and _3) were designed that commonly appear within the *TaSD-A1, TaSD-B1*, and *TaSD-D1* genes (Fig. 2a). Evaluation of the sgRNA design was performed using an *in vitro* Cas9 digestion assay. The Cas9 protein *in vitro*-assembled with the target_2 sgRNA exhibited complete digestion of the target genome sequence under the utilized conditions, while the target_1 and the target_3 sgRNAs were less efficient ((Supplementary Fig. 2).

**Figure 2.**
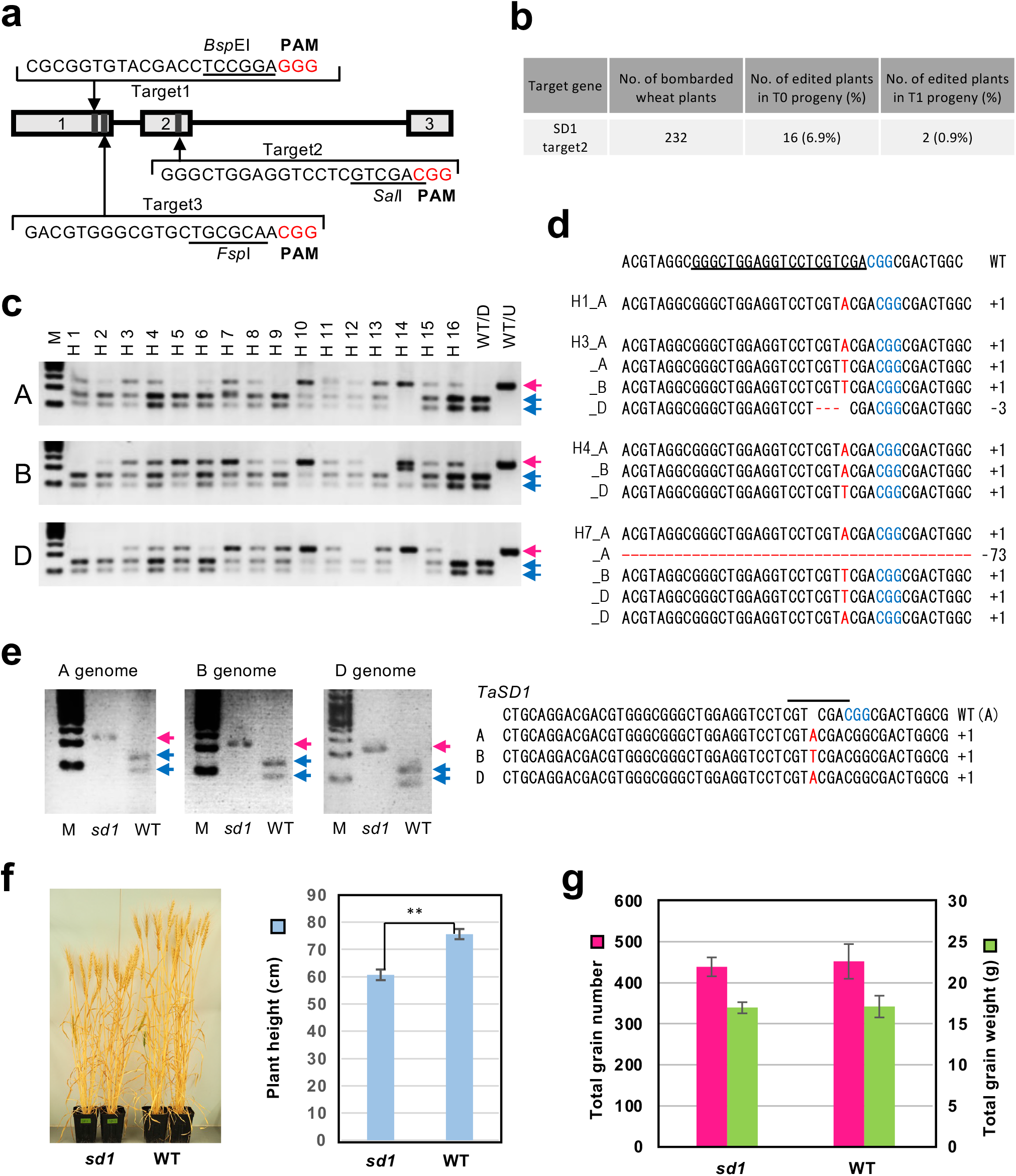
Introduction of *sd1* mutations in wheat. **a**, Target sequences conserved among the three homoeologous *TaSD1* genes were selected using the CRISPRdirect software. The locations of the target sequences are indicated by arrows. The boxes and lines indicate exons and introns, respectively. The three exons in *TaSD1* are numbered. **b**, Summary of the CAPS analysis of bombarded E_0_ plants and their progeny. **c**, CAPS assays of selected positive E_0_ plants using genome-specific primers. **d**, Mutations detected within the target region of positive E_0_ plants. The gRNA sequence is underlined in the WT sequence. Protospacer adjacent motif (PAM) sequences are indicated in blue letters. Insertions and deletions are indicated in red letters. **e**, A genome-specific CAPS assay of an *sd1* mutant line (H7-1, E_1_). The A, B, and D genome sequences of H7-1 are aligned with the A genome sequence of the WT. The inserted nucleotide and PAM sequence are indicated by red and blue letters, respectively. **f**, Comparison of plant stature of *tasd1* (left) and WT (right) plants. Average tiller height based on measurements of all plants. Data represent the mean ± SE of seven *sd1* and six WT plants. **g**, Comparison of grain yield. Average total grain numbers and average total grain weight for each plant are shown. The data represent the mean ± SE of seven *sd1* and six WT plants.

Gold particles coated with the CRISPR/Cas9 (target_2) RNP were bombarded into the SAMs of numerous imbibed wheat embryos, prepared as previously described, to enable large-scale screening for *tasd1* mutants. We observed undigested bands in 16 plants among the 232 bombarded embryos that had been grown into mature plants, representing 6.9% of the total bombarded embryos (Fig. 2b). A CAPS assay, using genome-specific primers, followed by Sanger sequence analysis of the undigested bands, revealed that the mutations were distributed among the A, B, and D genomes (Fig. 2c, Fig. 2d). Sixteen positively-selected E_0_ plants were subjected to E_1_ genotype analysis. The CAPS assay detected mutant alleles of *tasd1* genes in E_1_ plants derived from two E_0_ plants (H7 and H14, in Fig. 2c, Supplementary Fig.4). Among H7- and H14-derived E_1_ plants, the H7-1 plant did not display a digested band after *Sal* I treatment, suggesting that mutations had occurred in all six *TaSD1* genes (Supplementary Fig. 4, Fig.2c). The other E_1_ plants displayed digested bands, suggesting WT or partial mutations in the hexaploid genome. A CAPS assay with genome-specific primers indicated that the H7-1 E_1_ plant is a triple mutant (Fig. 2e). Sanger sequencing of the *Sal* I-resistant PCR amplicons revealed that the mutations in the H7-1 plant represent an A, a T, and an A insertion in the A, B, D genomes, respectively (Fig. 2e). These mutations caused frame shifts that resulted in putative mRNAs with a premature stop codon or no stop codon, suggesting that the TaSD1 function was knocked out in the H7-1 plant (Supplementary Fig. 6).

A primer set for *TaSD1* that spans an intron (common to the A, B, and D genomes) was designed and a semi-quantitative RT-PCR analysis was performed to analyze *TaSD1* expression in the H7-1 E_1_ plants. Results indicated that expression of the *TaSD1* genes was completely silenced in H7-1 E_1_ plants (Supplementary Fig. 5), suggesting the possibility of no-stop or nonsense-mediated mRNA decay.

The phenotype of the *tasd1* mutant was analyzed in the E_3_ generation of the H7-1 line of wheat plants. Both wild-type (WT) and H7-1 mutant plants were grown under long day conditions in an environmentally-controlled growth room. The mutant plants exhibited greener leaf color and shorter plant height. The average final height of the plants was approximately 10% lower in the *tasd1* mutant (Fig 2f), relative to the height of WT plants. The average total number of grains and grain weight were nearly equivalent in WT and *tasd1* plants (Fig. 2g).

We predicted potential off-target, candidate sites using Cas-OFFinder and identified 10 candidates having at least two mismatches in the site for target 2. Among them, eight candidates exhibited the same pattern: GGG<u>T</u>TGGAGGT<u>T</u>CTCGTCGAAGG (Underlined bases indicate the mismatches). Therefore, three candidates were selected from among the eight candidates, and five primer sets were designed (Supplementary Table 1). The amplicons produced from the five primer sets were subsequently sequenced and no mutations were found in the potential off-target regions. These data indicate that the mutations occurred without causing any off-target mutations.

In summary, we successfully applied genome editing on different gene loci with the iPB-RNP method utilizing wheat SAMs as a tissue source. In addition, we also created a wheat line carrying both *Rht-B1b* and *sd1* together using genome editing and demonstrated the cumulative effect of the two ‘Green Revolution’ semidwarf genes. The 10% reduction in plant height achieved would further contribute to lodging resistance in current, widely-used cultivars. RNP-based direct delivery of CRISPR/Cas9 has been successfully used to create genome-edited plants in several crop species^5,7,8^. The need for tissue/cell culture in the gene-editing methodology, however, hampers the broad utility of this approach for a wide range of commercial varieties in many crops, including maize and wheat. The iPB-RNP method described here represents an alternative approach for creating genome-edited wheat varieties. The efficiency of genome editing using the iPB-RNP method is comparable to the iPB-DNA method, which utilizes transient expression of CRISPR/Cas9 to accomplish genome editing^4^. Since no transgene integration occurs when using Cas9 RNPs, the application of iPB-RNP method in practical breeding and commercialization has the potential for broad impact to modern agricultural applications.

## Methods

### Preparation of SAMs

The protocol for SAM preparation has been previously described. In brief, mature seeds of wheat (*Triticum aestivum*) ‘Haruyokoi’ (*RhtB1-b, RhtD1-a*) were sterilized with sodium hypochlorite and imbibed at 25°C overnight. The coleoptile and the first three leaves, which cover the SAM, were removed from the embryo under a stereo microscope using an insulin pen needle 34G (ϕ0.2 mm; TERUMO, Japan). The embryos were separated from the endosperm and placed upright in a petri dish containing Murashige and Skoog (MS) basal medium supplemented with maltose (30 g/L), 2-morpholinoethanesulfonic acid (MES) monohydrate (0.98 g/L, pH 5.8), a plant preservative mixture (3%; Nacalai Tesque, Japan), and phytagel (7.0 g/L; Sigma Aldrich, USA). Thirty embryos were placed on the medium in each petri dish for each cycle of particle bombardment.

### Preparation of Cas9 protein and sgRNA

Recombinant *Streptococcus pyogenes* Cas9 protein was purified from *Escherichia coli* as previously described^16^. Single-stranded, guide RNA (sgRNA) was prepared using a GeneArt™ Precision gRNA synthesis Kit (Thermo Fisher Scientific, USA). The templates for the *in vitro* transcription were designed and amplified using appropriate primers (Supplementary Table 1, Supplementary Table 3) according to the manufacturer’s instructions.

### *In vitro* digestion of CRISPR/Cas9 RNP

DNA fragments containing the target sites were amplified from genomic wheat DNA using designated primer sets (Supplementary Table 1, Supplementary Table 3), purified and dissolved in RNase-free water. Cas9 protein (0.2 µg) and sgRNA (0.2 µg) were mixed and left for 10 min at room temperature to form an RNP complex. The RNP was incubated with the purified target DNA (100–200 ng) in CutSmart^®^ buffer (New England BioLabs) in a total volume of 10 μL, and digestion was allowed to proceed for 1 h at 37°C. The digested products were then separated on a 3% agarose gel.

### Preparation of microprojectiles and biolistic delivery

Gold particles coated with Cas9 RNP were prepared as previously described^7^ with slight modification. The purified Cas9 protein (12 μg) and sgRNA (5 μg) were mixed in a binding buffer (20 μL) containing 5 μL of 10×CutSmart^®^ buffer and 1μL of RNase inhibitor (40U, Takara, Japan) and left for 10 min at room temperature. After addition of 5 μL of TransIt transfection reagent (TaKaRa), the mixture was allowed to sit for an additional 5 min. Two hundred and seventy micrograms of gold particles (0.6 μm, InBio Gold, Australia) were added to the RNP mixture, tap-mixed, and then left to sit for 10 min. The gold particles were subsequently dispersed by slight sonication and 5 μL of the mixture was loaded onto a hydrophilic film (Scotchtint, 3M, Japan), placed on a macrocarrier and allowed to air-dry at room temperature for 15 min. Bombardment was conducted using a PDS-1000/He™ device (Bio-Rad, USA) with a target distance of 6.0 cm from the stopping plate. The vacuum in the chamber was 27 inches of Hg and the helium pressure was 1350 psi. Bombardment was repeated three times per plate.

### Cleaved amplified polymorphic sequences (CAPS) analysis

Genomic DNA was extracted from the fifth leaf of E_0_ progeny and the first leaf of the E_1_ progeny as previously described^4^ For *TaOr* (AK457010.1) and *HPGP-like* (AK333546.1), PCR amplification was conducted using KOD FX Neo DNA polymerase (Toyobo, Osaka, Japan) with gene specific primers (300 nM of each), and genomic DNA (50 ng). The mixture was denatured for 2 min at 98°C in a thermocycler and then subjected to 30 cycles of amplification (98°C for 30 s, 60°C for 30 s, 68°C for 30 s). For *TaQsd1* (LC209619.1) and *TaSD1*, PCR amplification was conducted using TaKaRa LA Taq^®^ with GC buffer (TaKaRa), gene specific primers (300 nM of each), and genomic DNA (50 ng). The mixture was denatured for 2 min at 94°C in a thermocycler and then subjected to 30 cycles of amplification (94°C for 30 s, 55°C for 30 s, 72°C for 20 s). The common primer and genome-specific primer sets used in the PCR are listed in Supplementary Table 1. The amplified PCR products were digested with *Pst* I (TaQsd1), *Nde* I (HRGP-like1_t2), *Sal* I (TaSD1_t2) or Cas9 RNPs (TaOr_t0 and TaOr_t1) and subsequently analyzed by agarose gel electrophoresis. Undigested bands from the restriction enzyme digestion were purified and cloned into the pGEM-T easy vector (Promega, USA) or using Zero Blunt™ TOPO™ PCR Cloning Kit (Invitrogen™, USA), and sequenced on a 3130xl Genetic analyzer (Applied Biosystems, USA).

### Plant growth conditions

Twelve hours after bombardment, the embryos were transferred to a basal MS medium and cultured for 2–3 weeks in a growth chamber under long day conditions (16 h light/8 h darkness, 25°C). The seedlings were subsequently planted in pots (3 seedlings/pot, ϕ10.5 cm) and grown in a phytotron under long day conditions (16 h light/8 h darkness, 20°C).

### RT-PCR analysis

Total RNA was isolated from leaf tissue using an RNeasy Mini Kit (Qiagen, Hilden, Germany) according to the manufacturer’s protocol. First-strand cDNA was synthesized from total RNA (0.5 μg) using a PrimeScript™ II 1st strand cDNA Synthesis Kit (TaKaRa, Japan). PCR was conducted with TaKaRa LA Taq^®^ with GC buffer (TaKaRa) as follows: initial denaturation (94°C for 1 min), followed by 28 cycles (98°C for 10 s, 55°C for 15 s, and 72°C for 30 s) using specific primers for 18s rRNA and *TaSD1* (Supplementary Table 1).

### Phylogenetic tree

The amino acid sequences of GA20ox proteins were obtained from the Gramene database (http://www.gramene.org/). A phylogenetic tree was constructed using the neighbor-joining method. Bootstrap values were calculated from 1000 replicates.

### Off-target detection

PCR analysis was conducted to detect off-target mutagenesis in E_1_ mutants. Off-target sites were identified by Cas-OFFinder (http://www.rgenome.net/cas-offinder/)^17^. DNA was isolated from the first leaf of E_1_ plants, as previously described^15^. Each PCR was conducted using TaKaRa LA Taq^®^ with GC buffer (TaKaRa), according to the manufacturer’s instructions, along with the designated primers (300 nM of each: Supplementary Table 1) and genomic DNA (50 ng). The mixture was denatured for 2 min at 94°C in a thermocycler and then subjected to 30 cycles of amplification (94°C for 30 s, 55°C for 30 s, 72°C for 30 s). The resulting PCR products were then sequenced and analyzed.

### Sequencing analysis

PCR products used in the CAPS analysis and off-target detection were cloned into pCR-BluntII-TOPO (Thermo Fisher Scientific, USA) and sequenced on a 3130xL genetic analyzer (Applied Biosystems, USA).

### Data Availability

All data generated or analyzed during this study are included in this published article (and its Supplementary Information files). Regarding sequence data, the NCBI GenBank identifiers are: LC209619.1 (*TaQsd1*), AK457010.1(*TaOr*), AK333546.1 (*TaHRGP-like*) and LN828667.1 (*TaSD1*).

## Acknowledgements

We are grateful to Dr. Yukiko Osawa for the help in CAPS screening of the mutants. We also thank Dr. Masaki Endo for helpful discussions. This work was supported in part by grants from JSPS KAKENHI (JP17K19249) and NARO Bio-oriented Technology Research Advancement Institution (Research program on development of innovative technology) to R.I.

## Author contributions

R.I. conceived and supervised the study. Y.K. and R.I. designed the experiments. Y.K., Y.L., W.L., J.Z. and M.K. conducted the experiments. W.L., J.Z., M.K., H.H., Y.N., N.T., and R.I. analyzed data. Y.K., W.L. and R.I. wrote the manuscript.

## References

1. Goldberg, R. B., Beals, T. P. & Sanders, P. M. Anther Development : Basic Principles and Practical Applications. Plant Cell 5, 1217–1229 (1993).

2. Reiser, L. & Fischer, R. L. The ovule and the embryo sac. Plant Cell 5, 1291– 1301 (1993).

3. Imai, R. et al. In planta particle bombardment (iPB): A new method for plant transformation and genome editing. Plant Biotechnol. 37, 171–176 (2020).

4. Hamada, H. et al. Biolistic-delivery-based transient CRISPR/Cas9 expression enables in planta genome editing in wheat. Sci. Rep. 8, 14422 (2018).

5. Woo, J. W. et al. DNA-free genome editing in plants with preassembled CRISPR-Cas9 ribonucleoproteins. Nat. Biotechnol. 33, 1162–1164 (2015).

6. Toda, E. et al. An efficient DNA- and selectable-marker-free genome-editing system using zygotes in rice. Nat. Plants 5, 363–368 (2019).

7. Svitashev, S., Schwartz, C., Lenderts, B., Young, J. K. & Mark Cigan, A. Genome editing in maize directed by CRISPR–Cas9 ribonucleoprotein complexes. Nat. Commun. 7, 13274 (2016).

8. Liang, Z. et al. Efficient DNA-free genome editing of bread wheat using CRISPR/Cas9 ribonucleoprotein complexes. Nat. Commun. 8, 1–5 (2017).

9. Liu, Y. et al. In planta Genome Editing in Commercial Wheat Varieties. Front. Plant Sci. 12, 648841 (2021).

10. Hedden, P. The genes of the Green Revolution. Trends Genet. 19, 5–9 (2003).

11. Peng, J. et al. ‘Green revolution’ genes encode mutant gibberellin response modulators. Nature 400, 256–261 (1999).

12. Monna, L. et al. Positional Cloning of Rice Semidwarfing Gene, sd-1 : Rice “Green Revolution Gene” Encodes a Mutant Enzyme Involved in Gibberellin Synthesis. 17, 11–17 (2002).

13. Sasaki, A. et al. Green revolution: A mutant gibberellin-synthesis gene in rice. Nature 416, 701–702 (2002).

14. Spielmeyer, W., Ellis, M. H. & Chandler, P. M. Semidwarf (sd-1), ‘green revolution’ rice, contains a defective gibberellin 20-oxidase gene. Proc. Natl. Acad. Sci. 99, 9043–9048 (2002).

15. Peng, J. et al. The Arabidopsis GAI gene defines a signaling pathway that negatively regulates gibberellin responses. Genes Dev. 11, 3194–3205 (1997).

16. Hamada, H. et al. An in planta biolistic method for stable wheat transformation. Sci. Rep. 7, 11443 (2017).

17. Kunitake, E. et al. CRISPR/Cas9-mediated gene replacement in the basidiomycetous yeast Pseudozyma antarctica. Fungal Genet. Biol. 130, 82–90 (2019).

18. Bae, S., Park, J. & Kim, J. S. Cas-OFFinder: A fast and versatile algorithm that searches for potential off-target sites of Cas9 RNA-guided endonucleases. Bioinformatics 30, 1473–1475 (2014).

