## Supplemental Figures and Tables for "*In planta genome* editing with CRISPR/Cas9 ribonucleoproteins"

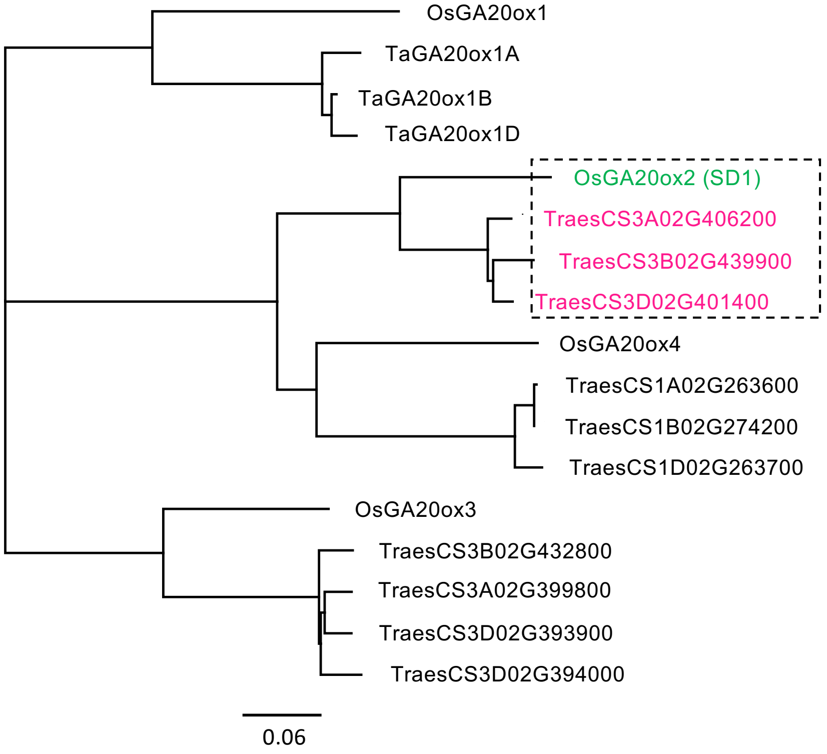


**Supplementary Fig 1. Phylogenetic tree of GA20ox from rice and wheat.** The amino acid sequences of GA20ox were obtained from the Gramene database (http://www.gramene.org/). A phylogenetic tree was constructed using the neighbor-joining method. The rice SD1 clade is boxed.

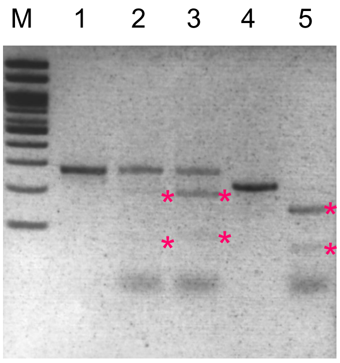


**Supplementary Figure 2. In vitro Cas9 cleavage analysis.** A genomic fragment containing target 1 and 3 was PCR-amplified (lane 1) and digested with Cas9/gRNAtarget1 (lane 2) and Cas9/gRNAtarget3 (lane 3), respectively. A genomic fragment containing target 2 (lane 4) was also digested with Cas9/gRNAtarget2 (lane 5). Stars denote digested bands.

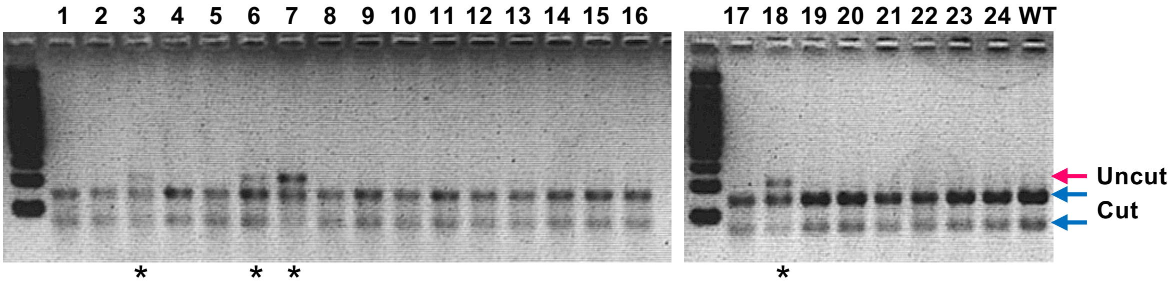


**Supplementary Figure 3. CAPS-based screening of sd1 mutations using tissue from the 5th leaf of bombarded T0 plants.** A portion of the cleaved, amplified polymorphic sequences (CAPS) assay data for T0 screening. Genomic DNA was isolated from the 5th leaf of the main culm of WT ('Haruyokoi') and bombarded T0 plants. A universal primer set (SD1 target 2F and SD1 target 2R) was used to amplify all three homoeologous genes. Red and blue arrows

indicate undigested and digested bands after *Sal* I treatment, respectively. Stars denote samples displaying positive signals.

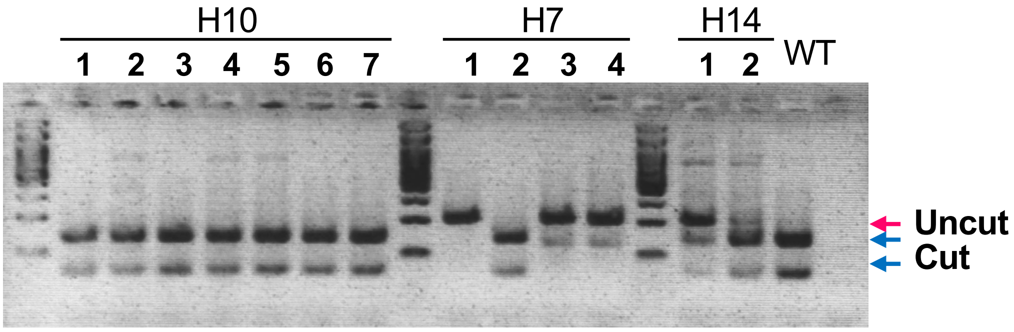


**Supplementary Figure 4**. **CAPS analysis of T1 plants**. Leaf tissue of $T_1$ plants from $T_0$ positive plants (H7, H10, H14) was utilized to conduct a CAPS analysis with *Sal* I digestion. Genomic DNA was isolated from the 1st leaf of WT and the $T_1$ plants. A universal primer set (SD1 target 2F and SD1 target 2R) was used to amplify all three homoeologous genes. Red and blue arrows indicate undigested and digested bands after *Sal* I treatment, respectively.

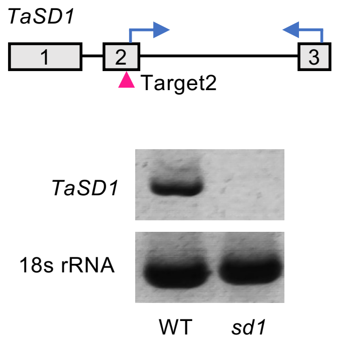


**Supplementary Figure 5. Semi-quantitative RT-PCR of SD1 expression in the *sd1* triple mutant (H7-1).** Primer locations are indicated by arrows. Targeted mutation site is indicated by red arrowhead. semi quantitative RT-PCR was carried out with leaf tissue of T1 plants.

```
WT-A:     MDTSPATPLLLQPPAPSIDPFAAKAAVNKNGGAATAVYDLRREPKIPAPFVWPHAEVRPTTAEELAVPVVDVGVLRNGDA
H7-1_A:   MDTSPATPLLLQPPAPSIDPFAAKAAVNKNGGAATAVYDLRREPKIPAPFVWPHAEVRPTTAEELAVPVVDVGVLRNGDA

          AGLRRAVAQVAAACATHGFFQVSGHGVDDALARAALDGASGFFGLPLAEKQRARRVPGTVSGYTSAHADRFASKLPWKET
          AGLRRAVAQVAAACATHGFFQVSGHGVDDALARAALDGASGFFGLPLAEKQRARRVPGTVSGYTSAHADRFASKLPWKET

          LSFGFHDRAGAAPVVVDYFTSTLGPDYEPMGRVYQEYCEKMKELSLRIMELLELGLGVEKRGYYRDFFADSSSIMRCNYY
          LSFGFHDRAGAAPVVVDYFTSTLGPDYEPMGRVYQEYCEKMKELSLRIMELLELGLGVEKRGYYRDFFADSSSIMRCNYY

          PPCPEPERTLGTGPHCDPTALTILLQDDVGGLEVLVDGDWRPVRPVPGAMVINIGDTFMALSNGRYKSCLHRAVVNRRQE
          PPCPEPERTLGTGPHCDPTALTILLQDDVGGLEVLVRRRLAARPPRPRRHGHQHRRHLHGAVERAVQELPAPGGGEPAAG

          RRSLAFFLCPREDRVVRPPPGLRSPRRYPDFTWADLMRFTQRHYRADTRTLDAFTQWFSSSTSPPPPAPAAQQAA*
          AAVAGLLPVPARGPRGAAAAGPEEPAAVPGLHLGRPHALHAAPLPRRHAHPRRLHPVVLLLHLAAAARPGGPAGGLIASP

          DPIDPRADSPRGSRRGIFVGTSPRARAPPSQVRWARAEWCPRGFPAPHHLPFLDAGSRSLLLALFVTTRHMHA


WT-B:     MVLQTAQQEPSLTRPPHCSAASARSPAAMDTSPATPLLLQPPAPSIDPFAAKAAVNKNGGAATAVYDLRREPKIPAPFVW
H7-1_B:   MVLQTAQQEPSLTRPPHCSVASARSPAAMDTSPTTPLLLQPPAPSIDPFAAKAAVNKNGGAATAVYDLRREPKIPAPFVW

          PHAEVRPTTAEELAVPVVDVGVLRNGDAAGLRRAVAQVAAACATHGFFQVSGHGVDDALARAALDGASGFFGLPLAEKQR
          PHAEVRPTTAEELAVPVVDVGVLRNGDAAGLRRAVAQVAAACATHGFFQVSGHGVDDALARAALDGASGFFGLPLAEKQR

          ARRVPGTVSGYTSAHADRFASKLPWKETLSFGFHDRAGAAPVVVDYFTSTLGPDYEPMGRVYQEYCEKMKELSLRIMELL
          ARRVPGTVSGYTSAHADRFASKLPWKETLSFGFHDRAGAAPVVVDYFTSTLGPDYEPMGRVYQEYCEKMKELSLRIMELL

          ELGLGVEKRGYYRDFFADSSSIMRCNYYPPCPEPERTLGTGPHCDPTALTILLQDDVGGLEVLVDGDWRPVRPVPGAMVI
          ELGLGVEKRGYYRDFFADSSSIMRCNYYPPCPEPERTLGTGPHCDPTALTILLQDDVGGLEVLVRRRLAPRPPRPRRHGH

          NIGDTFMALSNGRYKSCLHRAVVNRRQERRSLAFFLCPREDRVVRPPPGLRSPRRYPDFTWADLMRFTQRHYRADTRTLD
          QHRRHLHGSVERAVQELPAPRGGEPAAGAAVAGLLPVPARGPRGAAAAGAEEPAAVPGLHLGRPHALHAAPLPRRHAHPR

          AFTQWFSSSTSPPPPAPAAQQAA*
          RLHPVVLLLLLLLLLLGPGGGLILLLPIDPRADSTRGSRHEFLSGPAHVRAPPFSGAVARRGVVPTWISGPTPPSIFGRW

          LASPPPSLVCHDSPYACPLL


WT-D:     MDTSPATPLLLQPPAPSIDPFAAKAAVNKGGGAATAVYDLRREPKIPAPFVWPHAEVRPTTAQELAVPVVDVGVLRNGDA
H7-1_D:   MDTSPATPLLLQPPAPSIDPFAAKAAVNKGGGAATAVYDLRREPKIPAPFVWPHAEVRPTTAQELAVPVVDVGVLRNGDA

          AGLRRAVAQVAAACATHGFFQVSGHGVDEALARAALDGASGFFRLPLAEKQRARRVPGTVSGYTSAHADRFASKLPWKET
          AGLRRAVAQVAAACATHGFFQVSGHGVDEALARAALDGASGFFRLPLAEKQRARRVPGTVSGYTSAHADRFASKLPWKET

          LSFGFHDRAGAAPVVVDYFTSTLGPDYEPMGRVYQEYCGKMKELSLRIMELLELSQGVEKRGYYREFFADSSSIMRCNYY
          LSFGFHDRAGAAPVVVDYFTSTLGPDYEPMGRVYQEYCGKMKELSLRIMELLELSQGVEKRGYYREFFADSSSIMRCNYY

          PPCPEPERTLGTGPHCDPTALTILLQDDVGGLEVLVDGDWRPVRPVPGAMVINIGDTFMALSNGRYKSCLHRAVVNRRQE
          PPCPEPERTLGTGPHCDPTALTILLQDDVGGLEVLVRRRLAPRPPRPRRHGHQHRRHLHGNYSSLSVAFAD*

          RRSLAFFLCPREDRVVRPPPGLRSPRRYPDFTWADLMRFTQRHYRADTRTLDAFTQWFSSSSSSAQEAA*
```

**Supplementary Figure 6. Putative amino acid sequences of the mutant TaSD1 proteins.** The upper amino acid sequence is the wild type and the lower is the genome-edited amino acid sequence of TaSD1. Altered amino acids are indicated by red characters.

Supplementary Table 1. Sequences of the primers used in this study.

| Primer name | Primer sequence (5'-3') | Target | Purpose |
|---|---|---|---|
| TaQsd1 F | CAGCCTGGAGGGAATGACC | A, B and D genome | To amplify the TaQsd1 genome region target site |
| TaQsd1 R | ACCTGGTGGAATCCAGAGC | | |
| TaQsd1 gRNA F | TAATACGACTCACTATAGACGGATCCACCTCCCTG | TaQsd1 | Template DNA amplification for gRNA synthesis |
| TaQsd1 gRNA R | TTCTAGCTCTAAAACCTGCAGGGAGGTGGATCCGT | | |
| TaOr F | ACATCCATGGATCCTTTAGGATTAG | A, B and D genome | To amplify the TaOr genome region target (t0 and t1) sites |
| TaOr R | GCATAGTAGACCTTCAGATTAGCTG | | |
| TaOr t0 gRNA F | TAATACGACTCACTATAGGTAAGGGCCTACTAACC | TaOr_t0 | Template DNA amplification for gRNA synthesis |
| TaOr t0 gRNA R | TTCTAGCTCTAAAACCTGGTTAGTAGGCCCTTACC | | |
| TaOr t1 gRNA F | TAATACGACTCACTATAGTGGTAAGGGCCTACTAA | TaOr_t1 | Template DNA amplification for gRNA synthesis |
| TaOr t1 gRNA R | TTCTAGCTCTAAAACTGGTTAGTAGGCCCTTACCA | | |
| TaHRGPL1 F | AAGCTTCTGACTTCACTCAACAG | A, B and D genome | To amplify the TaHRGPL1 genome region target site |
| TaHRGPL1 R | TGAGGAAGTCTGCAACTCAG | | |
| TaHRGPL1 t2 gRNA F | TAATACGACTCACTATAGTTGGGCTCTCAGGGTAC | TaHRGPL1_t2 | Template DNA amplification for gRNA synthesis |
| TaHRGPL1 t2 gRNA R | TTCTAGCTCTAAAACTATGTACCCTGAGAGCCCAA | | |
| SD1 F | AGATGAAGGAGCTGTCGCTG | A and D genomes | To amplify the TaSD1 genome region target 1-3 sites |
| SD1 R | GGAAGCAGAGTGCAATCACG | | |
| SD1 F2 | GATCATGGAGCTGCTGGAGC | B genome | To amplify the TaSD1 target 1-3 sites |
| SD1 R2 | TGAAGGTGTCGCCGATGTTG | | |
| SD1 target 1:3 F | TCCCAGCATTGACCCGTTCG | Target 1 and 3 | To amplify the TaSD1 target1 and target3 sites for CAPS |
| SD1 target 1:3 R | ACACCTGGAAGAACCCGTGC | | |
| SD1 target 2 F | CGGACAGCAGCTCCATCATG | Target 2 | To amplify the TaSD1 target2 site for CAPS |
| SD1 target 2 R | GGTGTCGCCGATGTTGATGA | | |
| SD1 target 2A R | CTGATACGAGCAGCAGTAGC | Target 2A | Genome-specific primers for the TaSD1 target2 site amplification with the SD1 target 2 F |
| SD1 target 2B R | GCAAGGAAGGTGACCCAATT | Target 2B | |
| SD1 target 2D R | AGCAACGCTGAGAGAGGAGT | Target 2D | |
| gRNA target 1 F | TAATACGACTCACTATAGCGCGGTGTACGACCTCCGGA | Target 1 | Template DNA amplification for gRNA synthesis |
| gRNA target 1 R | TTCTAGCTCTAAAACTCCGGAGGTCGTACACCGCG | | |
| gRNA target 2 F | TAATACGACTCACTATAGGGCTGGAGGTCCTCGTCGA | Target 2 | Template DNA amplification for gRNA synthesis |
| gRNA target 2 R | TTCTAGCTCTAAAACTCGACGAGGACCTCCAGCC | | |
| gRNA target 3 F | TAATACGACTCACTATAGACGTGGGCGTGCTGCGCAA | Target 3 | Template DNA amplification for gRNA synthesis |
| gRNA target 3 R | TTCTAGCTCTAAAACTTGCGCAGCACGCCCACGT | | |
| SD1 RTPCR R | GCACGTGGGCTGGTCCCGACA | TaSD1 | RT-PCR |
| SD1 RTPCR F | TCATCAACATCGGCGACACC | | |
| Ta18s RTPCR F | GTGACGGGTGACGGAGAATT | 18s rRNA | RT-PCR |
| Ta18s RTPCR R | GACACTAATGCGCCCGGTAT | | |
| Off target 1 F | TCCCGCCTTTCTCTGAATG | 1026 | Off target detection |
| Off target 1 R | CCACCTTCACCAACACTTTC | | |
| Off target 2 F | TGCGCGATTCATCGGCCAAC | 1578 | Off target detection |
| Off target 2 R | AGTCTGGTGAACACCGCCAT | | |
| Off target 3 F | TTCCTAAGGACATTTGTGAGGTTA | 1642 | Off target detection |
| Off target 3 R | TCGTGGACATTGCTGCAACC | | |
| Off target 4 F | ACCAGCACCTTCTCCAGAAC | 1032 | Off target detection |
| Off target 4 R | CCTTGACGGAGACATTGGAC | | |
| Off target 5 F | CCCAGAATCTTCTCAGGGATAC | 2681 | Off target detection |
| Off target 5 R | ACTAGCACCTTCTCCAGAAC | | |

Supplementary Table 2. Analysis of possible off-target sites.

| # | Target | Chromosome Location | Position | Mismatch | Mutation |
|---|---|---|---|---|---|
| 1 | GGGCTGGAGGTtCTCGTCGgAGG | TGACv1_scaffold_014114_1AL | 1026 | 2 | – |
| 2 | GGGCTGGAGGTtgTCGTCGAAGG | TGACv1_scaffold_386600_5AL | 1578 | 2 | – |
| 3 | GGGtTGGAGGTtCTCGTCGAAGG | TGACv1_scaffold_117732_2AS | 1642 | 2 | – |
| 4 | GGGtTGGAGGTtCTCGTCGAAGG | TGACv1_scaffold_206147_3AL | 1032 | 2 | – |
| 5 | GGGtTGGAGGTtCTCGTCGAAGG | TGACv1_scaffold_660168_U | 2681 | 2 | – |
| 6 | GGGtTGGAGGTtCTCGTCGAAGG | TGACv1_scaffold_007802_1AL | 2718 | 2 | N.D. |
| 7 | GGGtTGGAGGTtCTCGTCGAAGG | TGACv1_scaffold_200959_3AL | 3374 | 2 | N.D. |
| 8 | GGGtTGGAGGTtCTCGTCGAAGG | TGACv1_scaffold_654903_U | 3265 | 2 | N.D. |
| 9 | GGGtTGGAGGTtCTCGTCGAAGG | TGACv1_scaffold_215894_3AS | 2609 | 2 | N.D. |
| 10 | GGGtTGGAGGTtCTCGTCGAAGG | TGACv1_scaffold_680557_U | 304 | 2 | N.D. |

N.D: Not determined. Red letters indicate mismatches

Supplementary Table 3. gRNA target sites.

| gRNA | Target (5'-3') | NCBI accession ID |
|---|---|---|
| TaQsd1 | ACGGATCCACCTCCCTGCAGCGG | LC209619.1 |
| TaOr_t0 | GGTAAGGGCCTACTAACCAGGGG | AK457010.1 |
| TaOr_t1 | TGGTAAGGGCCTACTAACCAGGG | AK457010.1 |
| TaHRGPL1_t2 | TTGGGCTCTCAGGGTACATATGG | AK333546.1 |
| TaSD1_t1 | CGCGGTGTACGACCTCCGGAGGG | LN828667.1 |
| TaSD1_t2 | GGGCTGGAGGTCCTCGTCGACGG | LN828667.1 |
| TaSD1_t3 | GACGTGGGCGTGCTGCGCAACGG | LN828667.1 |

The PAM motif in each target sequence is shown in red.
